# Peritubular macrophages phagocyte remains of undifferentiated spermatogonia in mouse testis

**DOI:** 10.1101/2022.10.14.512240

**Authors:** María Fernanda Marra, Jorge Ibañez, María Elisa Millán, Carlos Leandro Freites, Dario Fernandez, Luis Alberto Lopez

## Abstract

The cells involved in spermatogenesis are germ-cells, called spermatogonia, classified as: type A-undifferentiated, type A-intermediate and type B. During the spermatogenesis, more than 75% of the germ-cells undergo apoptosis and most of them are phagocyted by Sertoli cells. Peritubular macrophages in adult mouse testis are macrophages that both stimulate the proliferation and differentiation of undifferentiated spermatogonia in the wall of the seminiferous tubule. They have long processes and ramified appearance that squished between the lateral sides of neighbor myoid cells. We show, that a population of peritubular macrophages, grouped in pairs and activated, phagocyted undifferentiated spermatogonia in apoptosis. In adult mouse testis, 3.3x 10^5^ undifferentiated spermatogonia are in the germinal epithelium and 8,250 of them are in apoptosis. We counted in the testis 2,634.2 ±160 peritubular macrophages with phagocytic activity. If each one phagocyted one undifferentiated spermatogonia in apoptosis, it may indicated that peritubular macrophages phagocyted 31.9 % of the total undifferentiated spermatogonia in apoptosis. According to our knowledges, this is the first time that it is shown that undifferentiated spermatogonia in apoptosis are cleaned by peritubular macrophages.

**Summary Sentence:** We report that peritubular macrophages of adult mouse testis, phagocytic remains of apoptotic undifferentiated spermatogonia. These results show that peritubular macrophages, like Sertoli cells, participate in the remotion of germinal cells in the testis.

## Introduction

Spermatogenesis starts on the basal side of the testicular seminiferous tubules. The cells involved in this process are germ-cells, called spermatogonia. According to the amount of nuclear heterochromatin, these spermatogenic-cells are classified as type A, intermediate and type B. The most undifferentiated spermatogonia are called Asingle; it is a stem-cell which both renews itself by mitosis and divide by incomplete meiosis, forming a syncytial cyst of 2 (Apaired), 4 (Aaligned-4), 8 (Aaligned-8) or 16(Aaligned-16) spermatogonia. These cells represent the undifferentiated spermatogononia (Aundiff) that are found near the basement membrane of the seminiferous tubule and distributed among Sertoli cells. The differentiation of Aaligned spermatogonia generate A1(“differentianting”) spermatogonia, go through spermiogenesis, generating B (“differentiated spermatogonia”) that advance toward the tubule lumen (1–3).

During the spermatogenesis more than 75% of the germ-cells, undergoes apoptosis (4;5). Parallelly, the differentiation step of spermatogonia generates elongated spermatids which contain cytoplasmic portions whose must be eliminated as residual bodies (RB) (6). In both cases, the corresponding phagocytic elimination by Sertoli cells causes a reduction of the spatial competition, and a remotion of the harmful content released by the apoptotic germ cell necrosis (7). The Sertoli cell phagocytosis also removes autoantigens that can generate an autoimmune response.

The Sertoli cells engulf apoptotic germinal cells (AGC) and RBs by two main mechanisms. One is the interaction of type B scavenger receptor, presents in Sertoli cells, with phophatidylserine exposed on AGC and RBs (8–10). The another mechanism is mediated by the interaction of Tyro3, Axl, and Mer tyrosine kinase receptors (TAM receptors) with the growth arrest of gene 6. The extracellular N-terminal region of TAM receptors binds to the C-terminal domain of Gas6 resulting in activation of the intracellular tyrosine kinase domain and recognition of phosphatidylcholine expressed in the membrane of the AGC (11;12).

The seminiferous tubules (STs) possess a special microenvironment essential for spermatogenesis, generated for Sertoli cells that encompassing developing germ cells. The blood-testis barrier (BTB), that is formed by two adjacent Sertoli cells in the basal side of the ST is critical for maintaining the tissue homeostasis and immune microenvironment for normal germ cell development and protects the germinal cells of host’s immune systems. In this context, the Sertoli cells release immunoregulatory factors for the new antigens expressed by the spermatogonia on the cell surface, after that a systemic tolerance is established (13). Despite the aforementioned evidence, several studies question the fact that the BTB is considered the only structure for the protection of germ-cells, from the immune system (14). For example, in the early stage of the spermatogenesis, preleptotene spermatocytes and undifferentiated spermatogonia reside outside the BTB and produce antigenic proteins (15). In addition, certain germ-cell antigens behind BTB can escape into the interstitial spaces (14).

Macrophages are the main immune cells of mammals, and they are important for organogenesis, spermatogenesis, and male hormone production (16;17). In terms of morphology and location in the testis of adult-rodents, there are two well-characterized macrophage populations: the interstitial and peritubular macrophages. Interstitial macrophages contribute to maintain an immune-privileged environment in the organ (16;18), and they also play an important role in the development, regeneration, and production of testosterone (17; 19). Regarding to the peritubular macrophages, they represent a population of macrophages in the wall of the seminiferous tubule that secrete colony-stimulating factor and the enzymes involved in retinoic acid biosynthesis that both stimulate the proliferation and differentiation of undifferentiated spermatogonia (20). Peritubular macrophages, unlike interstitial ones, present on their surface the receptor for the major histocompatibility complex class II, suggesting that they are involved in the presentation of antigens (20).

In this work we showed that in the wall of STs of adult mouse testis, peritubular macrophages squish their cytoplasmic prolongations between adjacent myoid cells and engulf remains of undifferentiated spermatogonia in apoptosis. Also peritubular macrophages contain in its phagocytic compartment, phagolysosomes with remains of undifferentiated spermatogonia in apoptosis. These findings indicate that in addition to promoting the germ-cell differentiation in mouse testis, peritubular macrophages clearance undifferentiated spermatogonia in apoptosis.

### Animals, reagents and ST preparation

Male mice C-57/BL6 of three month old were born and maintained in our animal colony and housed in temperature and humidity-controlled rooms, with free access to water and food. Animals were maintained in accordance with the National Institutes of Health Guide for the Care and Use of Laboratory Animals. All procedures were approved by the Animal Research Committee of the Universidad Nacional de Cuyo, Argentina (CICUAL); Protocol 13/2012. At the time of sample collection, all animals were euthanized by carbon dioxide inhalation and efforts were made to minimize suffering.

ST of right testis from tree mice were removed and fixed with 4% paraformaldehyde or previously decapsulated and TS teased apart with needles and then fixed. The ST segments containing I-III, V-VI and VII-VIII stages (II-III, V-VI and VII-VIII ST segments in advance), were identified by transillumination in stereo microscopy (21) and dissected apart with a needle. The reagents used were from (Sigma Aldrich, St. Louis, USA) unless stated otherwise.

### Immunofluorescence staining

For thin section ST slides, testes were embedded in paraffin and 1 μm thickness sections were cut in a ultramicrotome (Ultracut R, Leica,Wien, Austria). The sections were deparaffinized, rehydrated, incubated with primary antibodies, washed with PBS and then incubated with secondary antibodies. Finally, the sections were washed and mounted with mounting solution: 0.2% Mowiol, 26% glycerol, 2-Phenylindole (DAPI) and 0.2 M Tris, pH 8.5).

For full-mount ST segments slides, fixed II-III, V-VI and VII-VIII ST segments, were washed with PBS, incubated with Ammonium Chloride to block auto-fluorescence and with primary antibodies (see Supplementary Table 1) in permeabilizing solution (PS): 0.05% saponin and 0.2% BSA in PBS), at RT, 12 h and with secondary antibodies in PS (Supplementary Table 2) at RT, 2 h, washed with a PBS solution and mounted with mounting solution, into a microscope slide camera of 0.3 μm high to avoid ST squeeze.

### Confocal microscopy

Thin section ST and full-mount ST segments slides were examined on an Olympus FV-1000 confocal microscope (Olympus America Inc., Center Valley, PA, USA) and the imagens processed with Fiji (Image J) software and edited with Adobe Photoshop 7.0 (Adobe Systems Inc., San José, CA, USA). Most of the figures shown in this work were z-stack from serial optical sections (OS) that were taken throughout the depth of the ST segment with step size of 400 nm generated from simple X,Y plane projections on the Z axis. The first OS was taken at the most superficial level of peritubular myoid cells (MC) and subsequent serial OS were taken until the deepest level of MC (near the germinal epithelium (22), Some figures were 3D reconstructed using Plagin 3D Viewer in mode surface and threshold for background filter.

### Area of peritubular wall of STs

The total surface area of peritubular wall of ST in the testis (ST wall area), was estimated considering to the ST as a cylinder where the surface is =*π.d.l. where **π*** is a constant, ***d*** the diameter of the ST cross-section and ***l*** is the total length of the STs. Samples (^S^) of segments of STs of know length (ST length^S^) were weighed (ST weight^S^) and the weight of total STs from a testis was calculated weighting the testis and discounting the weight of tunica albuginea and interstitial tissue (ST weight). Then, ST length = *ST length^S^. ST weight^S^/ST weight.* The ST length assayed was compensated 29 % due to the STs contraction by fixative procedure (23).

### Number of peritubular macrophages in the wall of STs

The number of peritubular macrophages (positive macrophages to Sox-3 ab) and peritubular macrophages containing VRUS in the cytoplasm (positive macrophages to both Sox-3 ab and E-Cadherin ab) by area of ST peritubular wall, were counted in 20 pictures of Z-stack images of II-III, V-VI and VII-VIII ST segments, from testes of each mouse, taken at 60X in a Olympus FV-1000 confocal microscope, using the cell counter tool of the Image J program (24). In each picture, an area (about of 197μm by 197μm, corresponding to 39,000 μm^2^ of ST area, were random selected. To estimate the total number of peritubular macrophages in the testis, the average number of macrophages in II-III, V-VI and VII-VIII ST segments per 39000 μm^2^, was extrapolated to the total area of ST surface.

### Statistical Analysis

The data are expressed as the average of three experiments ± SEM. Statistical significance was assessed with one-way analysis of variance (ANOVA). *P* ≥ 0.05 was considered significant.

## Results

### Peritubular macrophages engulfed remains of undifferentiated spermatogonia in the wall of ST

In immunohistochemistry-paraffin thin sections of mouse testis (Fig. 1A), peritubular macrophages stained with IBA-1 ab (25) localized in the wall of the ST, Fig 1B. Also, undifferentiated spermatogonia stained with SOX-3 ab (26) localized in both, in the germinate epithelium and in the wall of the ST, Figure 1C. It is interesting to observe a colocation of peritubular macrophages with undifferentiated spermatogonia in the wall of STs, Fig 1D.

**Figure 1.**
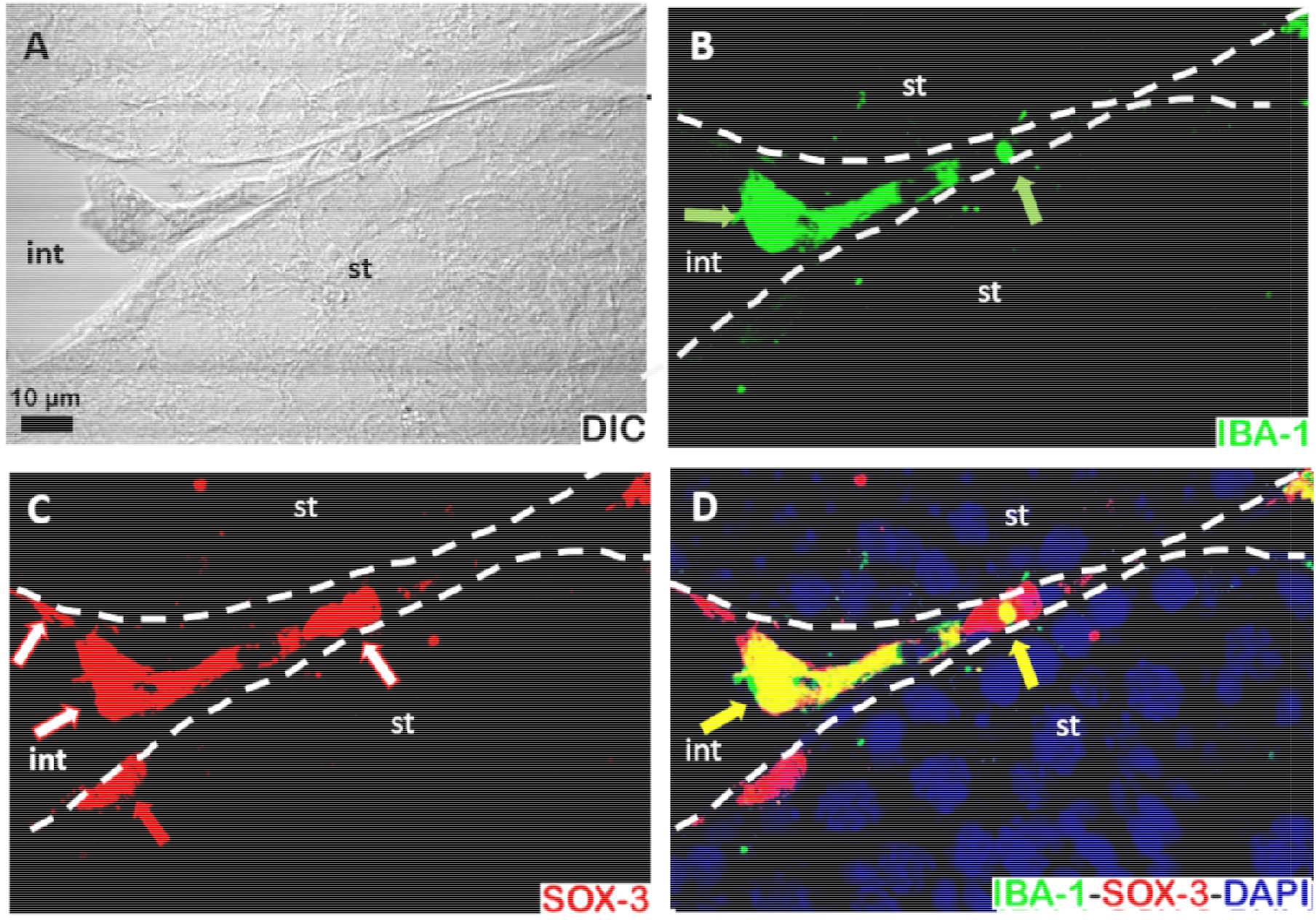
Peritubular macrophages and undifferentiated spermatogonia co-localize in the wall of seminiferous tubules. (A-D) Paraffin thin sections of mouse testis. Light field (A). Peritubular macrophages stained with IBA-1 ab, green and green arrows (B). Undifferentiated spermatogonia stained with SOX-3 ab in the peritubular wall, red and red arrows, and in germinal epithelium, red and white arrows (C). Merge of B and C and nucleous stained with DAPI (blue), co-localization of macrophages and undifferentiated spermatogonia in the wall of ST (yellow and yellow arrows) (D). Dashed lines indicate an outline of two STs throughout all figures; “st” refers to ST and “int” refers to interstitium throughout all figures. Scale bar, 10 μm

Using whole-mount of VII-VIII ST segment preparation, we observed a considerable number of peritubular macrophages (stained with IBA-1 ab) overlying to peritubular myoid cells (MC) in the peritubular wall (Figure 2A). We found 5.2, 5.8, and 7.9 peritubular macrophages per 39,000 μm^2^ ± SE (n=3) of peritubular wall area in II-III, V-VI and VII-VIII ST segments, respectively, where the number of peritubular macrophages was significantly major in VII-VIII ST segment (p≤ 0,05) (Table 1). Previous results shown that along of ST, VI-VII segment contain the highest density of peritubular macrophages and correlated with the highest density of undifferentiated spermatogonia in the underlying germinal epithelium (20). In this work we grouped in different way ST stages, but VII-VIII segment, with VII stage, contained the highest density of both peritubular macrophages and undifferentiated spermatogonia (Table 1).

**Figure 2.**
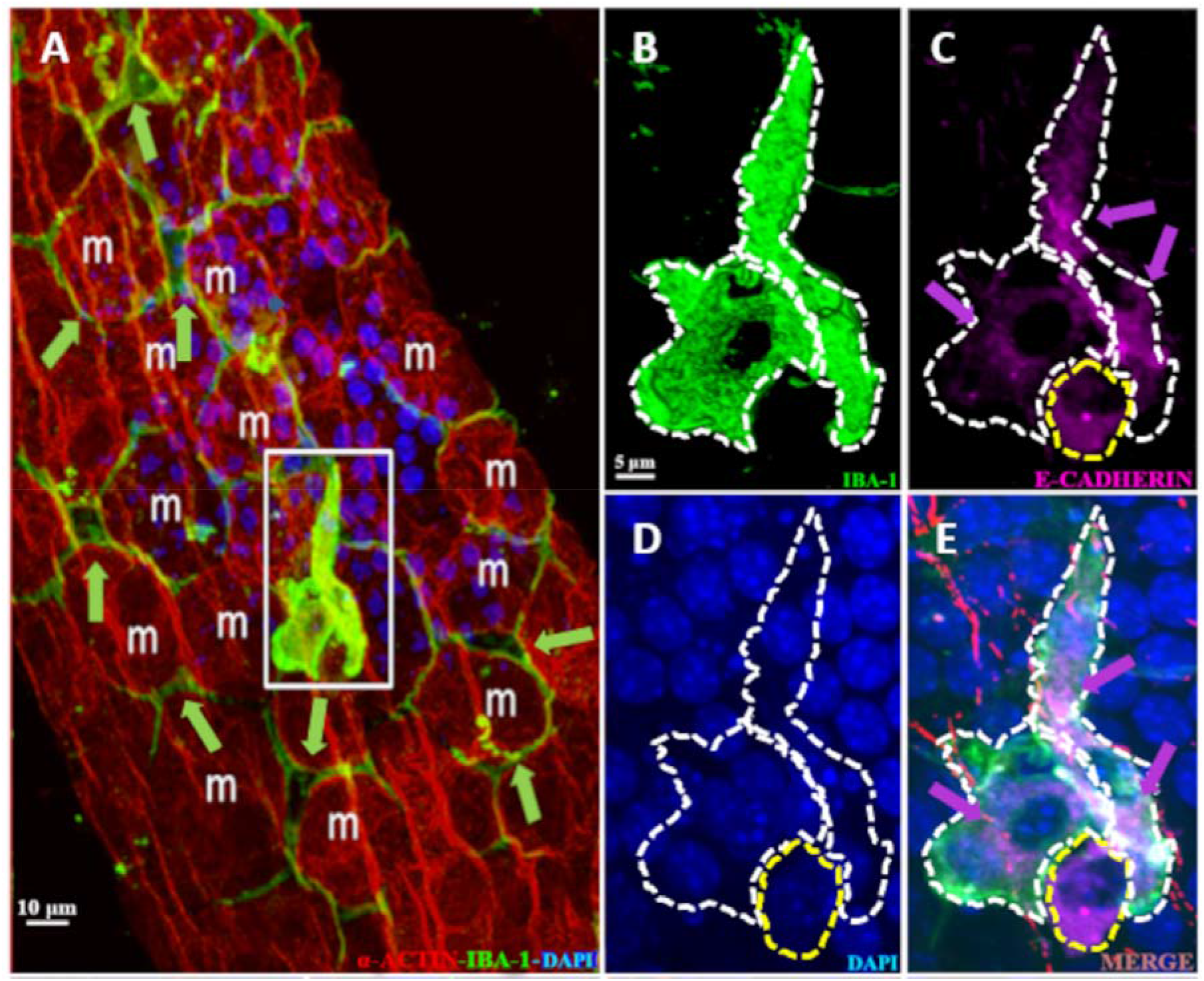
Peritubular macrophages with engulfed remains of undifferentiated spermatogonia in the wall of ST. (A-E) Immunofluorescent images of whole-mount of VII-VIII ST segment, where MC were stained with alpha-actin ab (red), peritubular macrophages with IBA-1 ab (green), undifferentiated spermatogonia and VRUS with E-Cadherin ab (magenta) and nucleous with DAPI (blue). Low magnification image, where MC are indicated with (“m”) and peritubular macrophages with geen arrows (A). High magnification of the boxed region in (A) of two macrophages (B-E), engulfed remains of undifferentiated spermatogonia (magenta) and VRUS (magenta and magenta arrows) (C) and nucleous (D). Merge of B-D, peritubular macrophages were made transparent to visualize engulfed remains of undifferentiated spermatogonia and VRUS thath are indicated with magenta arrows (E). Dashed white lines indicate an outline of two peritubular macrophages (B-E) and dashed yellow lines indicate an outline of engulfed remains of undifferentiated spermatogonia (C-E).

**Table 1 :** Number of peritubular macrophages in the wall of ST

| Number of peritubular macrophages | ST Segment | II-III<br>n±SE<br>(%±SE) | V-VI<br>n±SE<br>(%±SE) | VII-VIII<br>n±SE<br>(%±SE) | $\bar{X}$<br>n±SE<br>(% ±SE) |
| --- | --- | --- | --- | --- | --- |
| a:Total |  | 5.2±0,3<sup>a</sup> | 5.8± 0,1<sup>b</sup> | 7.9 ±0.1<sup>c</sup> | 6.3±0.6 |
| b:With engulfed remains |  | 0.46±0.12<br>(8.9±0.1) | 0.45±0.05<br>(7.8±0.2) | 0.44± 0.03<br>(5.6±0.1) | 0.45±0.01<br>7.40 ± 1.0 |
| c: With only VRUS |  | 0.7 ± 0.1<br>(13.5±1,2) | 0.6 ± 0.2<br>(10.3± 0.9) | 0.7± 0.1<br>(8.9±0.2) | 0.67±0.04<br>(10.9±1.4) |
| d:b+c |  | 1.16 ±0.2<br>(22.3±0.2 ) | 1.05±0.1<br>(18.1±0.1) | 1.14±0.3<br>(14.4±0.2) | 1.12±0.04<br>(18.3±2.3) |

(\*) n and % ± SE (n=3) of total (a), with engulfed remains (b) and sum of a+b (c) of peritubular macrophages/ 39,000 $\mu\text{m}^2$ of ST area in ST segments: II-III, V-VI, VII-VIII and average of the ST segments: $\bar{X}$ .(\*\*) <sup>c</sup> vs <sup>a,b</sup> p≤ 0,05

Most of the peritubular macrophages shown long processes and had ramified appearance that squished between the lateral sides of neighbor myoid cells Figure 2A and (20). As was previously mentioned, these branches did not penetrate the layer of germinal epithelium (see below) (20). We observed that same peritubular macrophages in the wall of ST, were grouped in pairs, looked more stained, presented higher body volume and contained less cytoplasm projections Figure 2A,B and E, and Supplementary Figure S1. According to previous reports, these morphologic characteristic correspond to activated macrophages (27). These activated peritubular macrophages engulfed cellular corpses, of about 15 μm diameter, with a small cytoplasm that surround nuclear material, from undifferentiated spermatogonia positive to E-Cadherin ab, (28) (Figure 2C and E and Supplementary Figure S1). We will mention in advance, to this population of activate peritubular macrophages, as peritubular macrophages with engulfed remains.

It is interesting to mention that the peritubular macrophages with engulfed remains also contained vesicles in the cytoplasm with remains of undifferentiated spermatogonia (stained with E-cadherin antibody) (these vesicles will be mentioned as VRUS in advance).The size of VRUS varied between 1 to 9 μm. See Figure 2C and E, Supplementary Movie S1. Supplementary Figure S2. Peritubular macrophages with engulfed remains always were detected in the wall of ST, Figure 2A,. Supplementary Figure S1, Supplementary Movie 1 and Supplementary Figure S2.

We asked whether the peritubular macrophages with engulfed remains where localized in particular segments of ST. We found that 0,46, 0,45 and 0,44 peritubular macrophages with engulfed remain per 39,000 μm^2^ of ST wall area in II-III, V-VI and VII-VIII segments respectively and represent 8.9, 7.8 and 5.6 % of total number of peritubular macrophages, respectively (Table 1), indicating that they did not localize to underlying particular areas of seminal epithelium.

### Same peritubular macrophages only contain VRUS

As we shown above, most of the activated peritubular macrophages with engulfed remain also contain VRUS, Figure 2C and E, Supplementary Movie 1 and Supplementary Figure S2.

Also, in a considerable number of peritubular macrophages, that does not contain engulfed remain, multiples VRUS was found in the cytoplasm, Figure 3A and D. We will name in advance these cells as peritubular macrophages with only VRUS. We found that 0.7, 0,6 and 0.7 peritubular macrophages with only VRUS per 39,000 μm^2^ of ST wall area were in the II-III, V-VI and VII-VIII segment respectively, indicating that they did not localize to underlying particular areas of seminal epithelium (Table 1).

**Figure 3.**
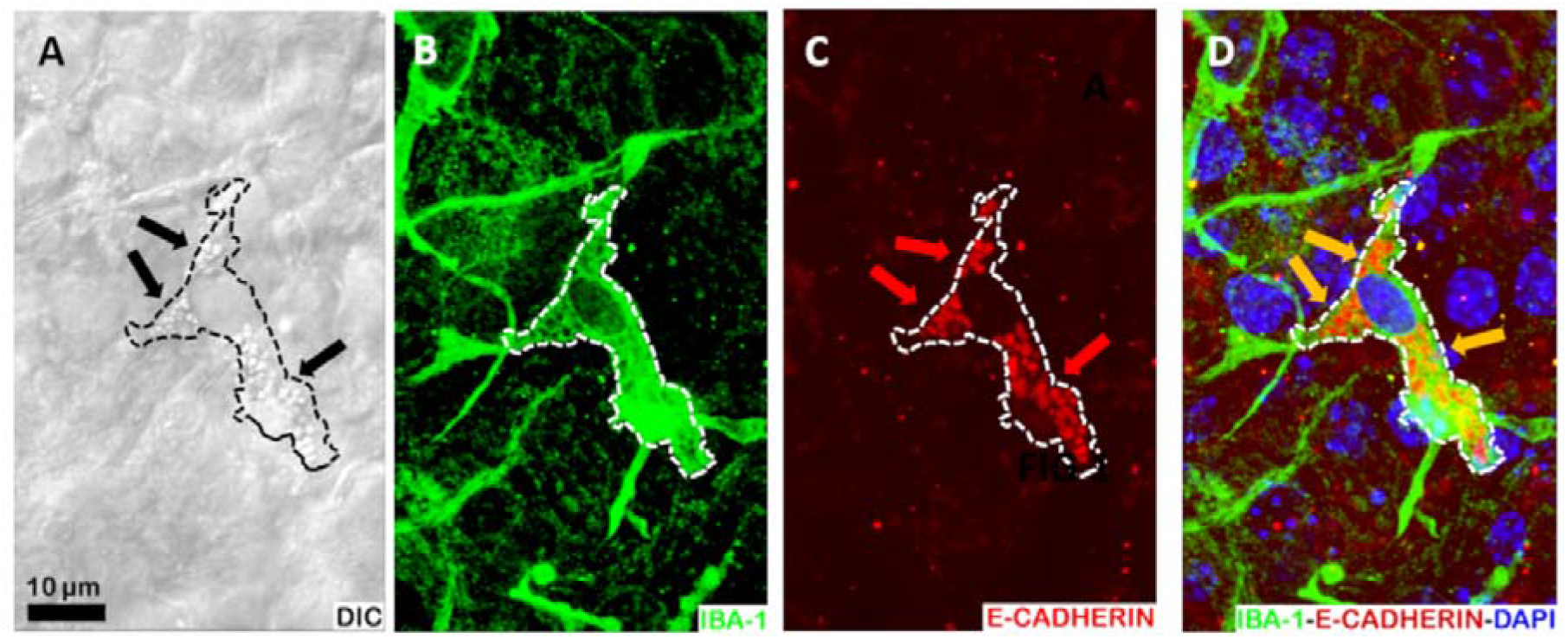
Some peritubular macrophages only contain VMUS. (A-D) Whole mount of V-VII ST segment, where peritubular macrophages and vesicles (black arrows) are visualized with light field (A). Peritubular macrophages stained with IBA-1 ab (green) (B), VRUS stained with E-Cadherin (reed and reed arrow) (C). Merge of B and C and nucleus stained with DAPI. Co-localization of VRUS with macrophage cytoplasm (yellow arrows) (D). Dashed lines indicate an outline of peritubular macrophages throughout all figures. Bar, 10 μm.

### VRUS in activated peritubular macrophages colocalized with phagolysosomes

To confirm whether remains of undifferentiated spermatogonia are in the phagocytic pathway of peritubular macrophages with only VRUS, we identify phagolysosomes in these cells. We used two markers of phagolysosomes, Cathepsin-D (lysosomal enzyme, (29)) and LAMP-1 (a protein associated with phagolysosomal membrane, (30)). We observed that Cathepsin-D colocalized with VRUS of peritubular macrophages with engulfed remains (Figure 4A-D). Also, LAMP-1 colocalized with VRUS of activated peritubular macrophages with only VRUS (Figure 4E-H). These results confirmed that undifferentiated spermatogonia remains has been phagocyted in peritubular macrophages.

**Figure 4.**
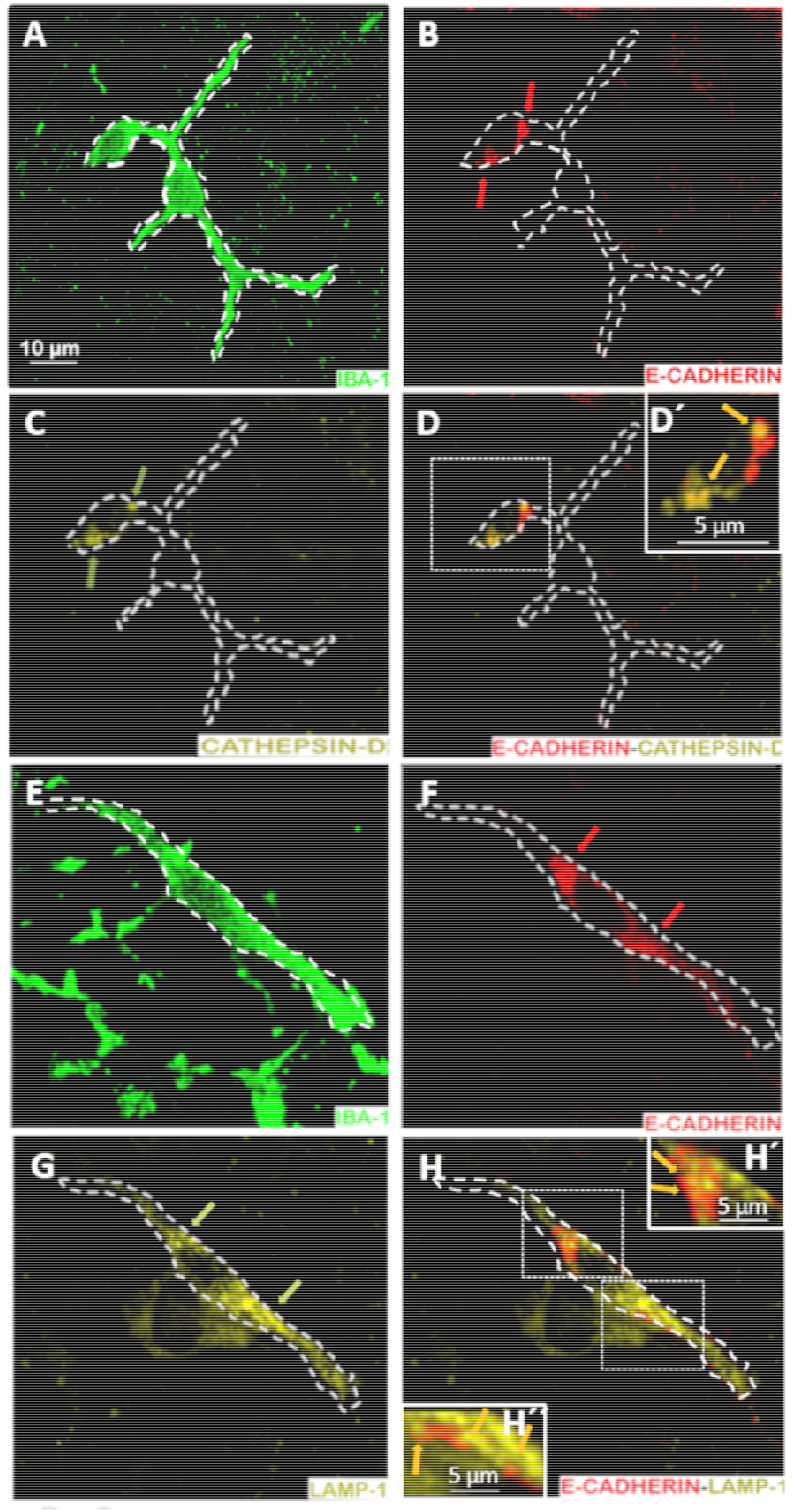
Peritubular macrophage contains remains of undifferentiated spermatogonia in the phagocytic pathway. **(A-D)** Peritubular macrophage in a whole mount of VII-VIII ST segment stained with anti-IBA-1 ab (green) (A), englobed remains of undifferentiated spermatogonia stained with E-cadherin ab (reed and reed arrows) (B), phagolysosomes stained with anti-Cathepsin-D ab (dark green and dark green arrows) (C). Merge of B and C (D), and high magnification of dashed boxed area where englobed remains co-localize with vesicles of phagolysosomes, magenta arrows (D’). Dashed white lines indicate an outline of peritubular macrophage throughout all figures. Bar, 10 μm and bar 5 μm in inserted high magnification imagen. (E-H) Peritubular macrophage in a whole mount seminiferous tubule stained with anti-IBA-1 Ab (green) (E), undifferentiated spermatogonia remains stained with anti-E-Cadherin Ab (reed and reed arrows) (F). Phagolysosomes stained with anti-LAMP-1 ab (yellow and yellow arrows) (G). Merge of F-G (H) and high magnification of two dashed boxed areas where englobed remains co-localize with vesicles of phagolysosomes, magenta arrows, up (H’) and down (H’’). Dashed white lines indicate an outline of peritubular macrophage throughout all figures. Bar, 10 μm and 5 μm in inserted high magnification imagens.

### Cellular corpses of undifferentiated spermatogonia come from remains of apoptotic cells

Due to the above results, that confirm that peritubular macrophages phagocyted remain of undifferentiated spermatogonia, we asked whether that remains come from apoptotic cells. Caspase 3 protein were used as signal of apoptosis activity (31), and we observed that remains of undifferentiated spermatogonia engulfed by peritubular macrophage were stained with anti-caspase 3 ab, Fig. 5 D-E.

**Figure 5.**
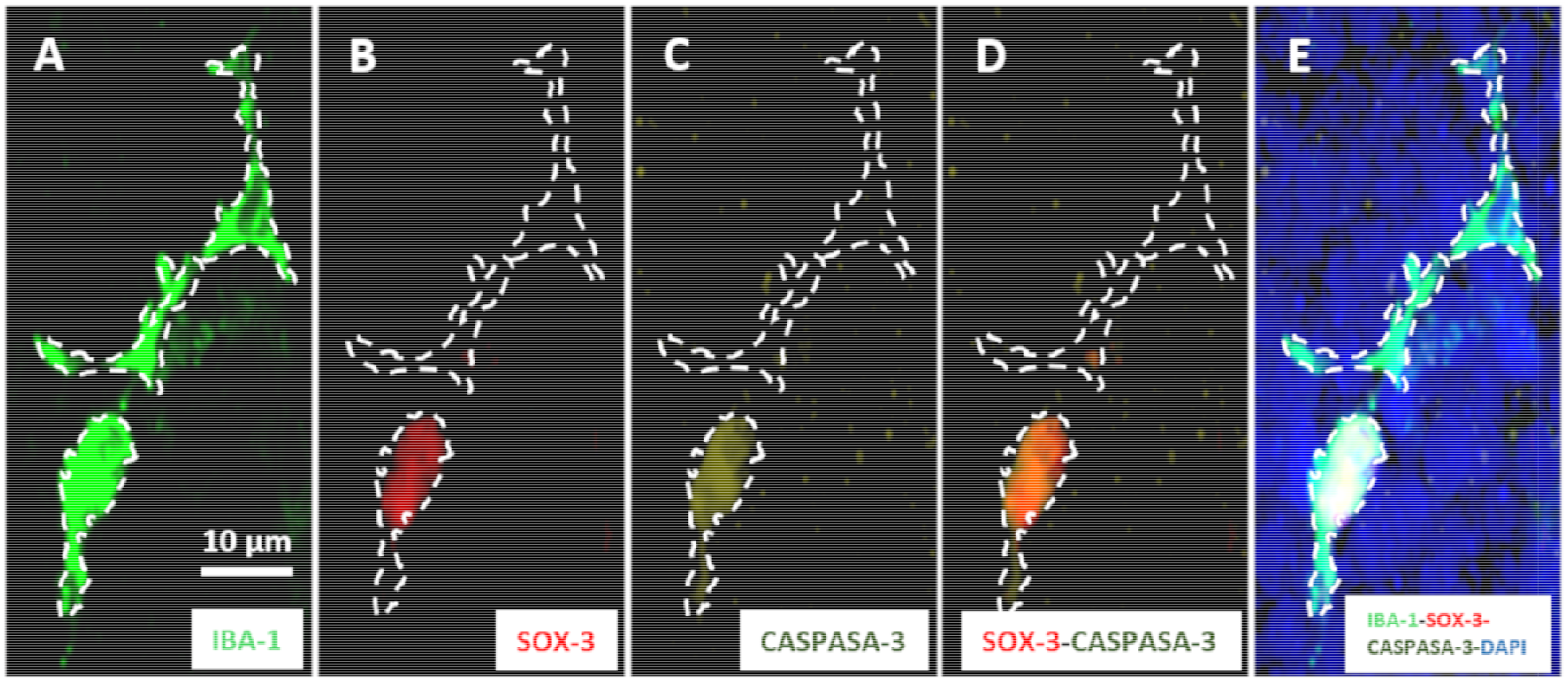
Remains of undifferentiated spermatogonia engulfed by peritubular macrophages were in apoptosis. (A-E) Immunofluorescent in a whole mount of VII-VIII ST segment, where peritubular macrophages were stained anti-Iba-1 ab, light green (A), cell corpse of undifferentiated spermatogonia stained with anti-SOX-3 ab, reed (B), cell corpse of undifferentiated spermatogonia stained with anti-caspase-3 ab, dark green (C). Merge of B and C, colocalization of reed and dark green, orange (D), and merge of A-D and DAPI (E). Dashed white lines indicate an outline of peritubular macrophages throughout A-E. Bar, 10 μm.

### A considerable number of peritubular macrophages phagocyted remains of undifferentiated spermatogonia

In the way to estimate the amount of peritubular macrophage with engulfed remains of undifferentiated macrophages in the ST wall area of the whole testis, we measured that the total length of STs in the testis was 1.9 ± 0.2 meter, similarly at the length reported previously (Nakata et al 2015), and that the total ST wall area of the testis was 8.6 ± 0,5 x 10^8^ μm^2^. Since that we detected an average of 6.3 peritubular macrophages in 39,000 μm^2^ (Table 1), in the total ST wall area of the testis there were 14,816 ± 1,740 peritubular macrophages. Considering that there were an average of 0.45 peritubular macrophages with engulfed remain in 39,000 μm^2^ of ST area (Table 1), in the whole testis there were 1,096 ± 97 peritubular macrophages with engulfed remains (7.4 % of total peritubular macrophages, Table 1). Meanwhile, there were 0.67 peritubular macrophages with only VRUS in 39,000 μm^2^ of ST area (Table 1), them in the whole testis there were 1,615.1 ± 153 of these cells (10.9 % of the total peritubular macrophages, Table 1). Considering the sum of the two population of peritubular macrophages, there were in the testis 2,634.2 ± 160 peritubular macrophages with phagocytic activity, that represent 18.3% of total peritubular macrophages in the testis, Table 1.

## Discussion

### Peritubular macrophages phagocyte undifferentiated spermatogonia in apoptosis

In the testis of adult mouse, peritubular macrophages are present in a considerable number along the segments of the ST ((20) and our results). We show in this paper that there were a number of 14,816 peritubular macrophages of which 18.3 % contain engulfed remains of undifferentiated spermatogonia (Table 1) and presents two characteristic, 7.4 % of total peritubular macrophages are peritubular macrophages with engulfed remains and 10.9 % were peritubular macrophages with only VRUS. One interpretation of these results is that peritubular macrophages population with engulfed remains, in the moment of obtain the samples, were engulfing cellular corpse of undifferentiated spermatogonia in apoptosis. Meanwhile, another peritubular macrophages population with only VRUS in the phagocytic pathway indicated than in the moment of obtain the sample, the engulfed corpse had been degraded and only remains VRUS. Another possibility is that peritubular macrophages with only VRUS, had phagocyted small size VRUS (see discussion below).

Considering that in adult mouse testis, 3.3x 10^5^ undifferentiated spermatogonia are in the germinal epithelium, that 8,250 (2.5 %) of them are in apoptosis (32), that there are 2,634.2 ±160 peritubular macrophages with phagocytic activity, that each one internalized remain of one undifferentiated spermatogonia, it may indicated that peritubular macrophages phagocyted 31.9 % of the total undifferentiated spermatogonia in apoptosis. According to our knowledges, this is the first time that it is shown that a considerable number of peritubular macrophages in apoptosis in the testis of adult mice are cleaned by peritubular macrophages.

### How peritubular macrophage, in the wall of ST, reached to remains of undifferentiated spermatogonia?

As was shown in Figure 2, in Supplementary Movie S1 and Supplementary Figure S2, and previously described by (20), peritubular macrophages are localized in the wall of ST, and their cytoplasm projection did not reach to the germinal epithelium, then, the phagocytosis of remains of undifferentiated spermatogonia by peritubular macrophages should occur in the wall of the ST.

It is known that in adult’s mouse testis, undifferentiated spermatogonia are localized in the base of germinal epithelium, behind the BTB that limits the transit of cells to the base of germinal epithelium (20), that apoptotic cells generate multiple signals including both “found” and “eat” me that conduct that they can be found and to be eaten by macrophages (33;34). But the behavior of peritubular macrophages and undifferentiated spermatogonia in apoptosis, mentioned above, did not explain how apoptotic remains of undifferentiated spermatogonia, come out of the germinal epithelium.

Apoptotic cells have a considerable capability to move into the tissues and also to pass to neighboring ones through basement membrane (34–36). The fact that undifferentiated spermatogonia are weak bound one to another in the germinal epithelium and do not need to cross the BTB to come up to the interstice, it suggest that migration of undifferentiated spermatogonia in apoptosis, would generated the encounter of undifferentiated spermatogonia in apoptosis with peritubular macrophage in the wall of the ST. But, according to our acknowledgments, penetration of peritubular macrophages to the germinal epithelium neither exits of intact or corpse of undifferentiated spermatogonia to the wall of seminiferous epithelium has not been reported.

In the male reproductive tract, spermatogonia release vesicles by exocytosis, known as extracellular vesicles (SVs) that participate in intercellular communication (37). SVs have three main sizes: *i.* Apoptotic bodies of more than 1000 nm diameter, than are usually produced by blebbling of the apoptotic cells; *ii.* Microvesicles with sizes between from 100 to 1000 nm than are formed by budding and shedding of the plasma membrane and *iii.* Exosomes, with sizes between 50 and 100 nm that are formed when multiple vesicles bodies fuse with the plasma membrane (37). It has been demonstrated that in the ST of adult’s mouse testis, spermatogonia secrets exosomes that has been found near the basemental membrane (38). Also, in mouse ST, it has been demonstrated that microvesicles and exosomes travel through the BTB to the basement membrane (35). Then, apoptotic bodies and other SVs from undifferentiated spermatogonia in apoptosis, in the germinal epithelium, could arrive to the wall of ST and be phagocyted by peritubular macrophages.

It has been demonstrated that macrophages attract apoptotic cells by inducing directional migration (39). Then it could be that peritubular macrophages tether dying undifferentiated spermatogonia close to basement membrane by use of CD14 (40) or by β_2_ integrin that could bind to complement fragment iC3b (41).

### Anti-inflamatory action of peritubular macrophages

It is necessary to consider, that in the testis, interstitial macrophages contribute to maintain an immune-privileged environment in the organ (16;18),

Peritubular macrophages, unlike interstitial ones, present on their surface the receptor for the major histocompatibility complex class II, suggesting that they are involved in the presentation of antigens (20).

The uptake of apoptotic cells usually suppresses the secretion from activated macrophages, of proimflammatory responses, preventing the secretion from activated macrophages of pro-inflammatory mediator such as tumor necrosis factor-α (6;42;43). Receptor-triggered release of the anti-inflammatory and immunosuppressive cytokine transforming growth factor-β1 by macrophages ingesting apoptotic cells might be crucial in mediating the autocrine or paracrine suppression of peritubular macrophage-directed inflammation (42–44). Rat peritubular macrophages, in activated phenotype in vitro, produce the anti-inflammatory interleukina-10, and impaired ability to stimulated T cell activation, in accordance with the role in maintaining testicular immune privilege (45). Peritubular macrophages can behavior as inactivated by the ligation of macrophage receptors mediating the engulfment of apoptotic cells, notably CD36 (43), its ligand thrombospondin and receptor for phosphatidylserine exposed by apoptotic cell. (46). Indeed, receptor-triggered release of the anti-inflammatory and immunosuppressive cytokine transforming-β1 by peritubular macrophages ingesting apoptotic cells might be crucial in mediating the autocrine or paracrine suppression of macrophage-directed inflammation (6;46).

### Peritubular macrophage phagocytosis could regular spermatogonia proliferation

Peritubular macrophages that are associated with regions of the ST containing undifferentiated spermatogonia, participate in multiples niches in the wall of the seminiferous tubule by secreting colony-stimulating factor and the enzymes involved in retinoic acid biosynthesis that both stimulate the proliferation and differentiation of undifferentiated spermatogonia (20). Nevertheless, peritubular macrophages with phagocytic activity do not localize to a particular region of the ST (Table 1). Recently, (47) has demonstrate in rat, that the exposure of peripuberal male rat to the phthalate metabolite mon-(2-ethylhexyl) phthalate (MEHP) cause testicular inflammation, spermatocyte apoptosis, and disruption of the BTB. MEHP treatment increased peritubular macrophage presence in the surface of STs and remained elevate by 2 weeks after exposure. Simultaneously, an increase of differentiated spermatogonia occurred 2 weeks after MEHP exposure. The peritubular macrophages phagocytosis of undifferentiated spermatogonia in apoptosis might bring signals to regulate undifferentiated spermatogonia to proliferate and differentiated.

## Author’ Contributions

L.A.L.and LF conceptualized the study. M.F.M. D.F. and M.E.M. designed, performed, and analyzed all experiments, with assistance from J.I.

The manuscript and figures were prepared by L.A.L., M.F.M. and JI

Funding was acquired by L.A.L. All authors reviewed the manuscript and approved the final version of the manuscript.

## Acknowledgment

The authors would like to thank to Estela Muñoz for help with antibodies.

## Abbreviations

ab: antibody
ST(s): seminiferous tubule(s)
BTB: bloot testis barrier
OS(s): Optical section(s)
MC: peritubular myoid cells
VRUS: cytoplasm vesicles with remains of undifferentiated spermatogonia stained with E-cadherin antibody
SV(s): extracellular vesicle(s)
RB: residual bodies
AGC: apoptotic germinal cells

