## supplementary Fig S1 for "Peritubular macrophages phagocyte remains of undifferentiated spermatogonia in mouse testis"

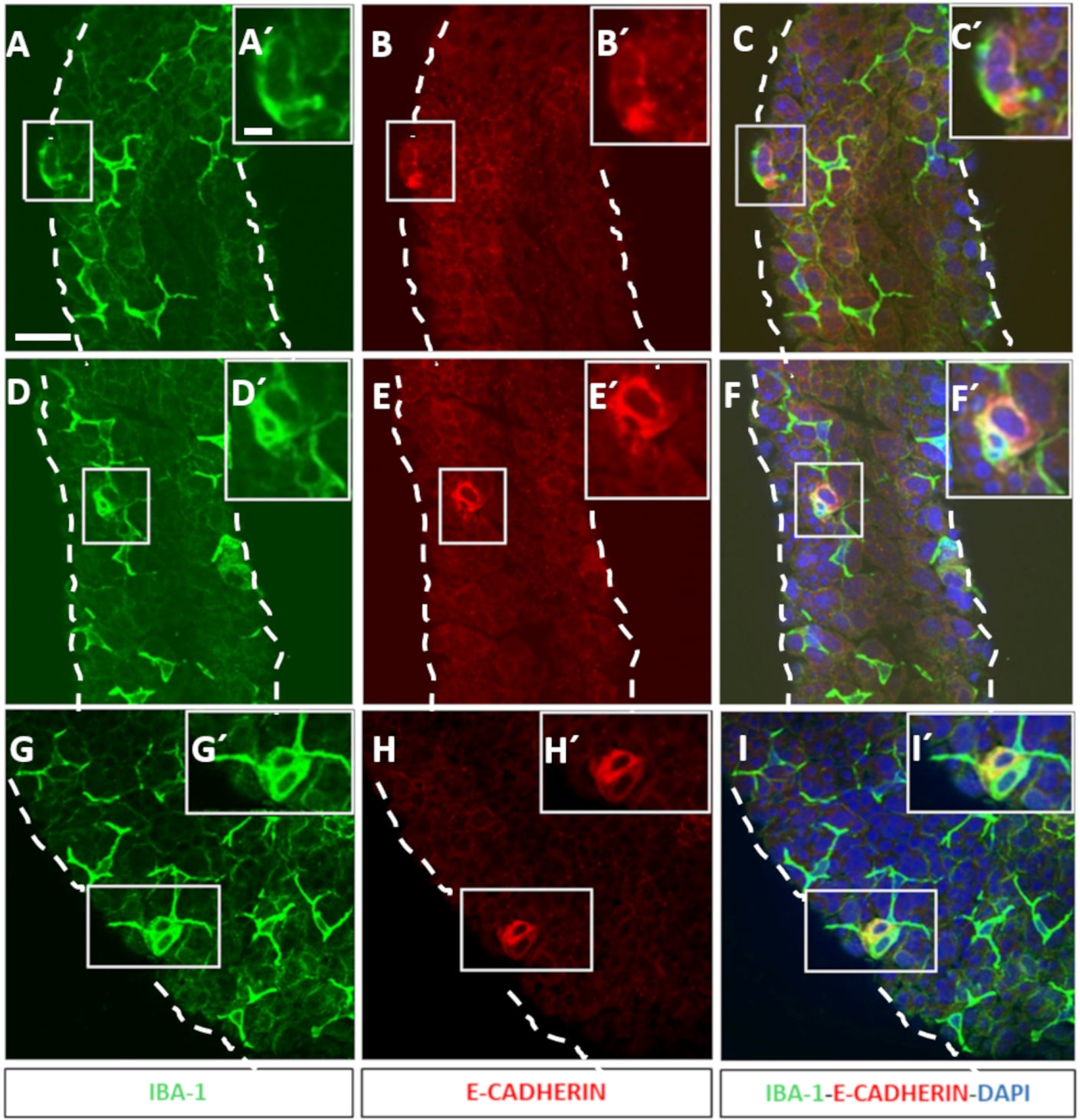

**Supplementary Figure S1. Peritubular macrophages engulfed remains of undifferentiated spermatozoa.** (A-I) Gallery of low magnification of immunofluorescent images of whole-mount of VII-VIII ST segment, where peritubular macrophages were stain with IBA-1 ab, green (A,D and G), undifferentiated spermatogonia with E-Cadherin ab, reed (B,E and H). Merge of A and B and nucleus stained with DAPI (C), D and E and nucleus stained with DAPI (F) and G and H and nucleus stained with DAPI (I). (A´-I´) Gallery of high magnification of the boxed region of peritubular macrophages A,D and G (A´,D´and G´), undifferentiated spermatogonia B,E and H (B´,E´and H´), and merge C,F and I (C´,F´ and I)´. Bars 40 µm
