## supplementary Fig S2 for "Peritubular macrophages phagocyte remains of undifferentiated spermatogonia in mouse testis"

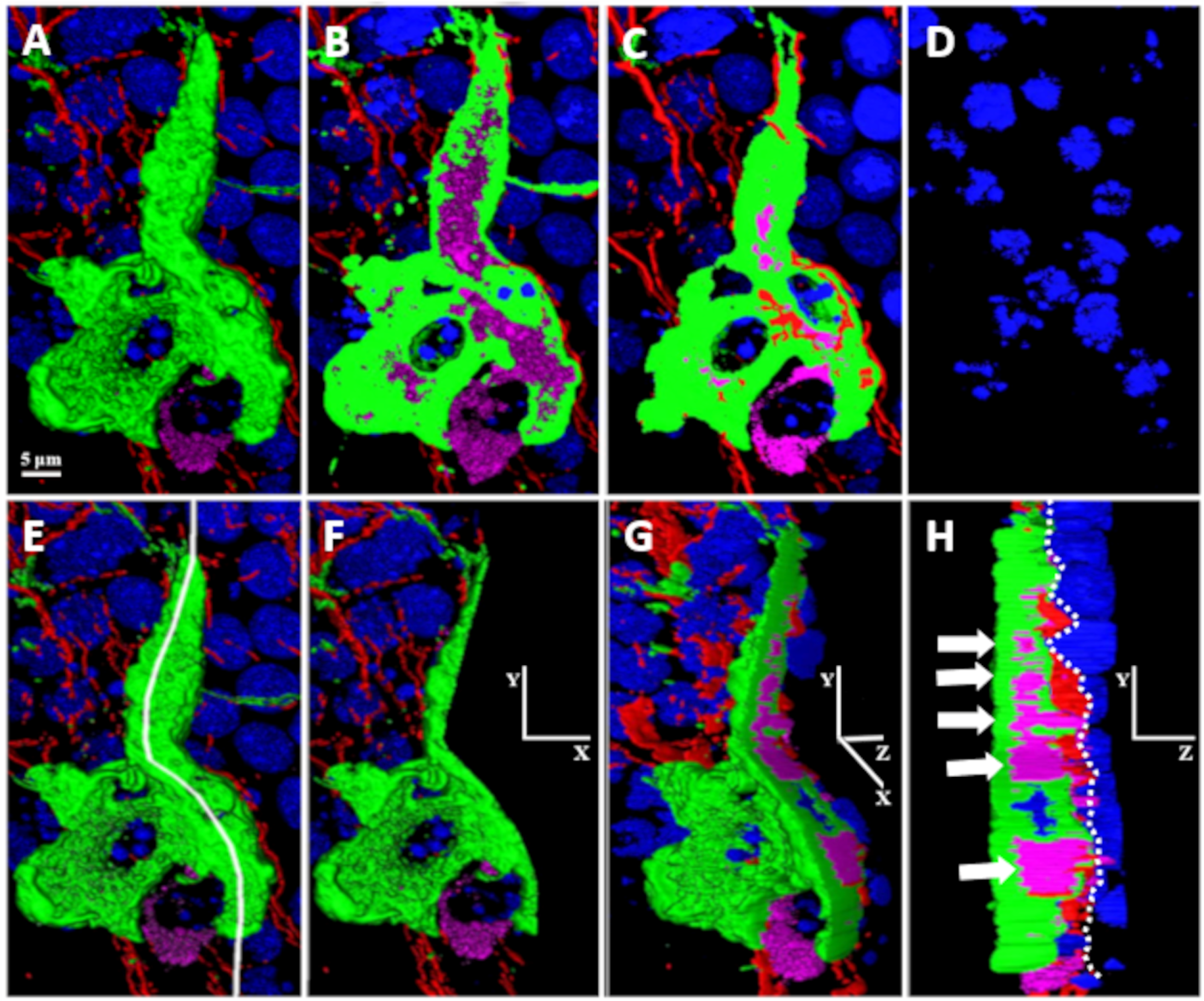

**Supplementary Figure S2. Selected OSs of Supplementary movie S1**. (A-D). OSs of planes X,Y with 1.1 µm in Z at: 0 µm (A), 5.5 µm (B), 9.6 µm (C) and 18 µm (D). (E-H). Reconstruction of A-D where a white line indicates the optical separation in two half (E) and left half of E (F). Rotation of F in YZ, 45 º (G) and 90ª (H). Dashed white linea in H indicates the limit between peritubular wall and seminal epithelium of ST (H). The white arrows indicate position of VRUS (H).
