## supplementary Tables S1 and S2 for "Peritubular macrophages phagocyte remains of undifferentiated spermatogonia in mouse testis"

**SUPPLEMENTARY MATERIAL**

**Supplementary Table S1: Primary antibodies used in immunofluorescence.**

| **Target** | **Supplier** | **Catalog number** | **Dilution**  **IF** | **RRID** |
| --- | --- | --- | --- | --- |
| **Alfa-actin** | Sigma-Aldrich | ab 5228 | 1/500 | AB_262054 |
| **Iba-1** | Abcam | ab 5076 | 1/100 | AB_2224402 |
| **E-cadherin** | Cell Signaling | ab 3195S | 1/200 | AB_2291471 |
| **Sox-3** | Santa Cruz | sc 101155 | 1/50 | AB_2195961 |
| **Lamp-1** | Cell Signaling | ab 15665S | 1/1000 | AB_2798750 |
| **Cathepsin-D** | Cell Signaling | ab 2284S | 1/200 | AB_10694258 |
| **E-Cadherin** | Santa Cruz | sc 8426 | 1/100 | AB_626780 |

**Supplementary Table S2: Secondary antibodies used in immunofluorescence.**

| **Target** | **Supplier** | **Catalog number** | **Dilution**  **IF** | **RRID** |
| --- | --- | --- | --- | --- |
| rabbit Alexa Fluor 488 | Jackson InmunoResearch | 711-545152 | 1/500 | AB_2313584 |
| goat Alexa Fluor 647 | Jackson InmunoResearch | 705-605-147 | 1/500 | AB_2340437 |
| goat Alexa Fluor 488 | Jackson InmunoResearch | 705-545147 | 1/500 | AB_2336933 |
| mause Cy3 | Jackson InmunoResearch | 715-165-151 | 1/500 | AB_2315777 |
